# Tuning plant promoters using a simple split luciferase method for quantifying transcription factor binding affinity

**DOI:** 10.1101/2023.02.13.528283

**Authors:** Y-M Cai, S Witham, NJ Patron

## Abstract

Sequence features, including the binding affinity of binding motifs for their cognate transcription factors, are important contributors to promoter behavior. The ability to predictably recode affinity enables the development of synthetic promoters with varying levels of response to known cellular signals. Here we describe a luminescence-based microplate assay for comparing the interactions of transcription factors with short DNA probes. We then demonstrate how this data can be used to design synthetic plant promoters of varying strengths that respond to the same transcription.

## Introduction

To initiate the transcription of synthetic genetic circuits, regulatory elements with predictable characteristics that respond to cellular signals are highly desirable. Similarly, the ability to rationally edit the sequence of endogenous sequences to alter the amplitude or timing of gene expression is useful for tuning quantitative phenotypes. Promoters play a role in initiating transcription through the recruitment of proteins, including transcription factors (TFs). While the availability of some proteins may influence expression patterns and levels^1^, the sequence features of promoters that encode the binding locations and affinity for TFs are critical for defining function^2^. In previous work, we developed computationally designed minimal synthetic promoters (MinSyns) for plants with binding sites for known classes of endogenous TFs^3^. We reasoned that manipulating binding sites to alter affinity would enable us to produce variants of individual MinSyns that respond to the same endogenous signal. To do this rationally, we required a method to investigate the impact of sequence variations in transcription factor binding motifs (TFBMs) on TF association.

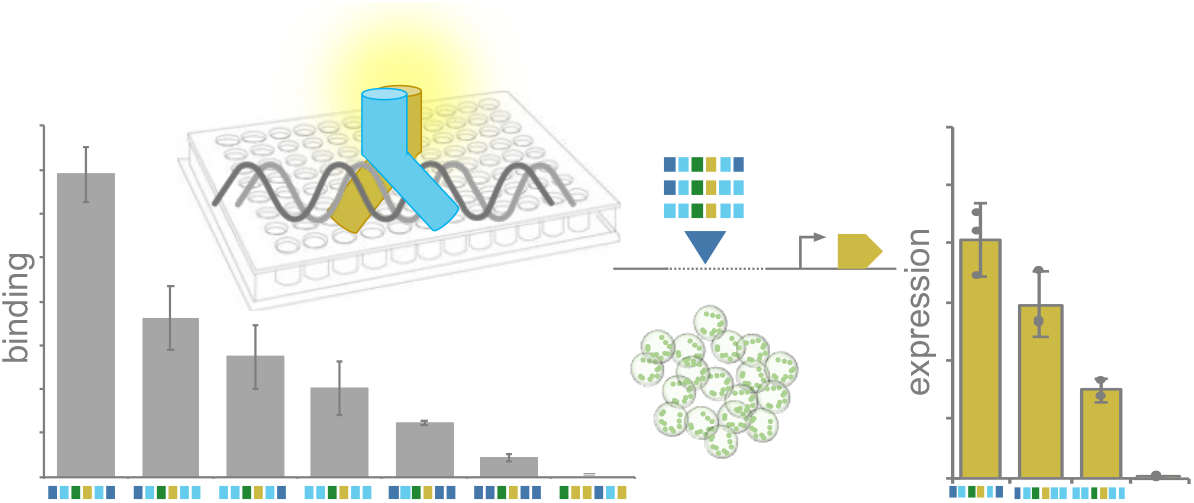

Several methods exist for assessing the binding of TFs to DNA. Electrophoretic Mobility Shift Assays (EMSA), in which chemically labeled DNA probes are combined with purified protein and protein-DNA complexes are identified by electrophoresis, are widely used but can be challenging to quantify and are labor-intensive. Surface plasmon resonance (SPR) has the advantage of generating real-time data but requires expensive specialized equipment. Systematic evolution of ligands by exponential enrichment (SELEX) and protein binding microarrays (PBM) offer high throughput approaches but, as a result, though cost per probe is low, have high total experiment costs. Fluorescent or colorimetric microplate-based protein-DNA affinity assays require no specialized equipment and are reasonably scalable and quantifiable^4^. However, labeling of DNA and the conjugation of chemical groups can add substantial time and cost. Here, we describe a luminescence-based microplate assay for the relative quantification of TF interactions with DNA probes (qTFD). This method uses short, unlabeled DNA probes, which are cheaply obtained, and minimal quantities of recombinant protein with a genetically encoded minimal (11 amino acid) HiBiT Tag. We demonstrate the that this assay can detect associations between several classes of TFs and probes with known target sequences. We then apply knowledge of TF-DNA interactions to modulate the strength of minimal synthetic plant promoters (MinSyns).

## Results and Discussion

### A low-cost assay for relative quantification of protein-DNA interactions

To avoid the use of expensive, surface-modified microplates and labeled DNA probes, we employed a commercially available DNA coating solution previously used for chemiluminescence immunoassay of DNA adducts^5^. This is used to bind short (60-80 bp) double-stranded probes to the plate. The amount of immobilized probe (F_DNA_) bound to the plate is assessed using PicoGreen. To enable the detection of protein, we expressed recombinant TF proteins with a C-terminal HiBiT tag^6^. Luciferase activity proportional to the level of HiBiT-tagged protein (L_TF_) is enabled by the addition of the LgBiT polypeptide. The amount of protein bound to the DNA probe is expressed as L_TF_/F_DNA_. To enable the comparison of this value to different probes, this is value is normalized to the value obtained for the same TF to a random sequence (Figure 1A). This provides a relative quantity (rQ) of bound protein in arbitrary units and differentiates between affinity for specific motifs and non-specific affinity for DNA.

**Figure 1:**
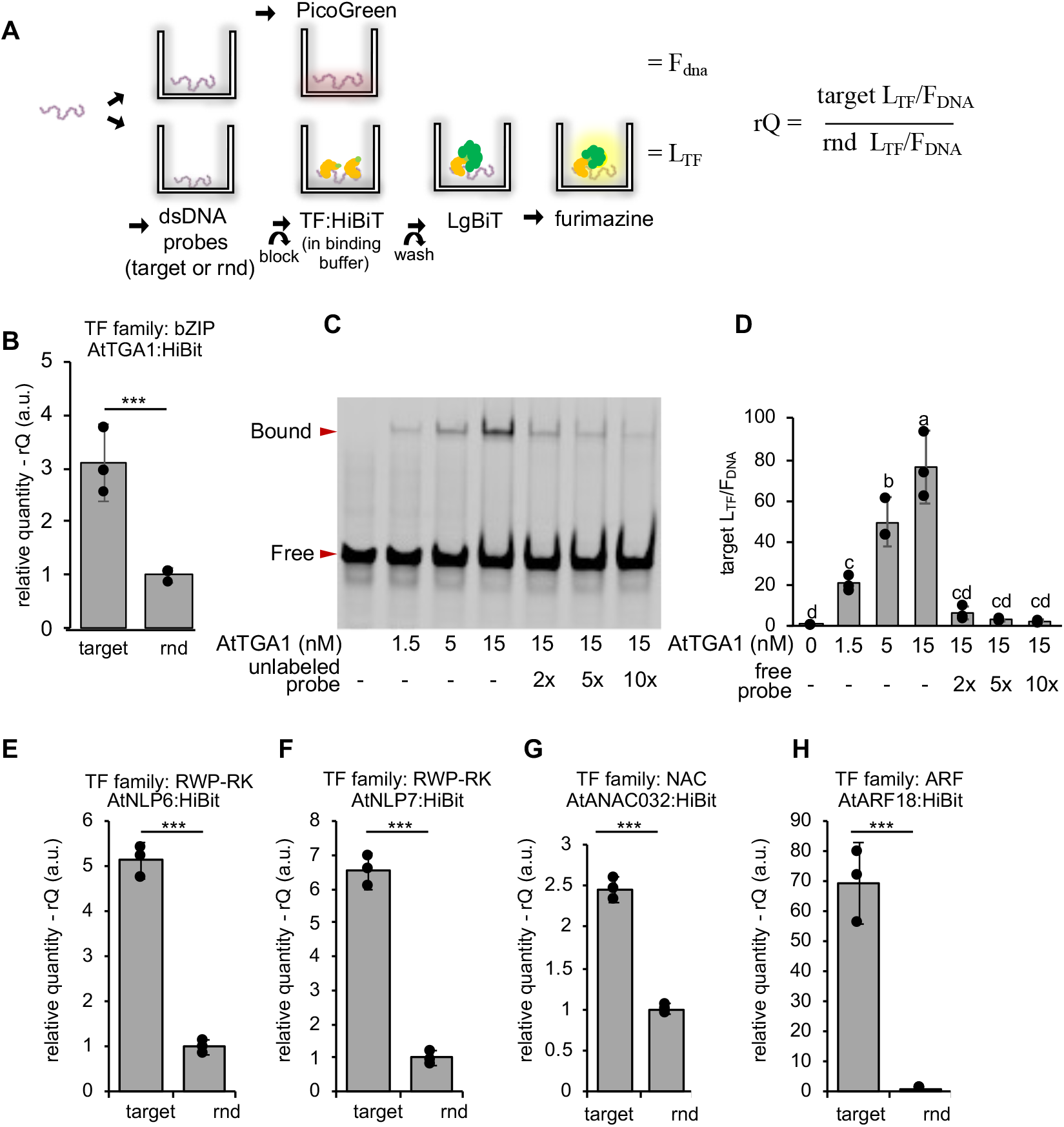
A low-cost assay for quantifying protein-DNA binding A. Schematic of the method for relative quantification of transcription-factor DNA binding (qTFD). DNA probes are added to six wells (three replicates for protein binding and three for quantifying immobilized DNA). Purified TF:HiBiT proteins are added to the former and, following removal of unbound protein, luminescence (L_TF_) is quantified by addition of the LgBit polypeptide and fumarizine substrate. Immobilized DNA is quantified with PicoGreen (F_DNA_). The amount of protein bound to the DNA probe is expressed as L_TF_/F_DNA_ and the relative quantity (rQ) is calculated by normalizing to values obtained for random (rnd) DNA. **B**. Detection of binding between AtTGA1 and DNA probes containing the as1 motif from the *CaMV35s* promoter compared to random DNA probes (rnd). **C**. An electromobility shift assay (EMSA) showing binding of AtTGA1 to an 80 bp labeled probe. The presence of an unlabeled competitor probe reduces binding to the labeled probe. **D**. Quantification of AtTGA1:HiBit-binding to the same probe using qTFD. The inclusion of unbound competitor probe in the binding buffer reduces binding to the immobilized probe. Values are the mean and 2 x standard error of 3 replicates. Different letters indicate significant differences (TukeyHSD). **E-H**. Detection of binding between four plant transcription factors to probes containing previously reported binding sites compared to random DNA probes (rnd). Values are the mean and 2 x standard error of 3 replicates. P-values were calculated using a t-test: ***, *P* ≤ 0.001.

We first exemplified the assay using *Arabidopsis thaliana* TGA1 (AtTGA1), which has previously been shown to bind to the as1 motif in the widely used *CaMV35S* promoter ^7^. First, we verified that this assay was able to detect AtTGA1-binding to probes with this motif (Figure 1B). We then benchmarked our assay to the electromobility shift assay (EMSA). In EMSA, the intensity of the protein-bound band increased with protein concentration from 1.5 to 15 nM (Figure 1B). The amount of protein detected in the qTFD assay (L_TF_/F_DNA_) mirrored this data (Figure 1C). Further, the inclusion of unlabeled (EMSA) or unbound (qTFD) competitor probes reduced binding (Figure 1B and C). We then demonstrated that qTFD works across TF families by demonstrating that we are able to detect significant binding (relative to random DNA) to probes containing previously reported targets of four TFs from the RWP-RK (or nodule inception (NIN)-like) family^8^, NAC (NAM, ATAF and CUC) family^9^ and ARF (auxin response factor) families (Figure 1D-G)^10^.

### Tuning plant promoters by relative quantification of TF-DNA interactions

Previously, we demonstrated the function of plant minimal synthetic promoters (MinSyns) containing three tandem pairs of a TGA1 TF binding site^3^. To tune expression of this promoter, we used publicly available data to identify six *Arabidopsis thaliana* genes for which a change in expression has been detected in direct response to nuclear localization of AtTGA1^11,12^. We used the FIMO software^13^ to identify candidate AtTGA1 binding sites within these target genes (Figure 2A) and, when we compared these using qTFD, found variations in binding (Figure 2B). TGA proteins are known to bind to DNA as dimers and, importantly, the binding of such dimers has been shown to be stabilized by the presence of other TGA homodimers at the site^14^. We therefore hypothesized that combinations of sites to which TGA proteins binding weakly and strongly could be used to tune the activity of promoters activated by AtTGA1. We selected two sites with strong and weak rQ (TFBS 02 and 12) and, maintaining the architecture of the TGA1-binding sites in our existing MinSyn^3^, we constructed promoters with different combinations (Figure 2C). These promoters were fused to a luciferase reporter gene (LucF) and normalized expression was determined following transfection in Arabidopsis mesophyll protoplasts. Three copies of the strong element gave high activity while expression from MinSyns with three copies of the weak element was non-significant. Interestingly, MinSyns with one strong but two weak elements also had significant activity, with expression correlating with the relative position of the strong site to the TSS (Figure 2D). This is consistent with the observation that TGA-DNA complexes can stabilize nearby dimers. In summary, we developed a luciferase-based microplate TF-DNA binding assay. We demonstrate the utility of this method for tuning expression of minimal synthetic plant promoters that respond to the same TF. We note that, as described, this assay does not provide equilibrium dissociation constants (KD) and that the arbitrary numerical values obtained for two different TFs to the same probe are not directly comparable. The ability to characterize sequence variants that influence TF binding will also be useful for informing the rational editing of endogenous genes to tune expression patterns and levels.

**Figure 2:**
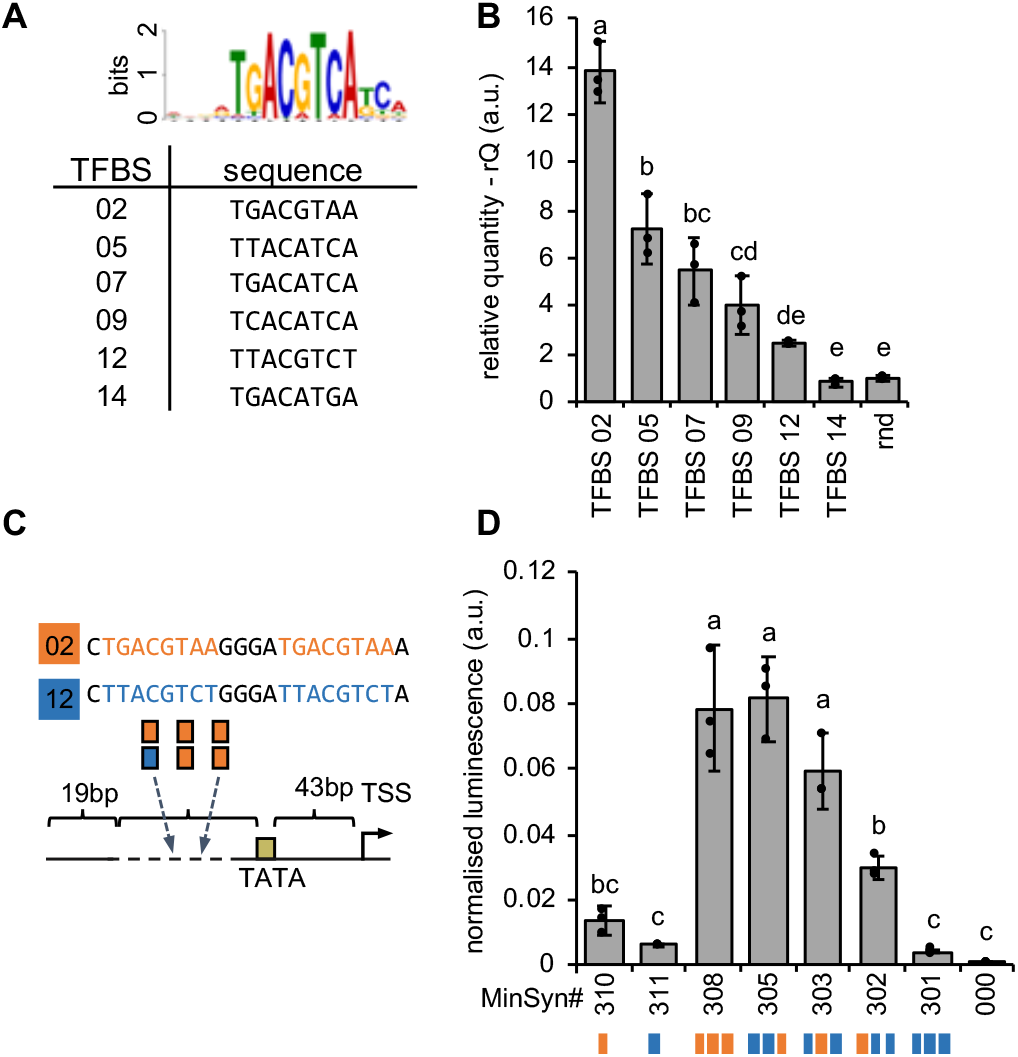
Tuning synthetic promoters by modulating binding sites. **A.** The position weight matrix consensus logo for AtTGA1 (above) and candidate binding sites from six Arabidopsis promoters that change expression in response to AtTGA1. **B**. Relative binding of AtTGA1:HiBiT to probes containing AtTGA1 binding sites. Error bars indicate the mean and 2 x standard error of 3 replicates. Different lowercase letters indicate significant difference (ANOVA, *P* = 2e-16, TukeyHSD, α= 0.05). C. Three pairs of AtTGA1 binding sites are inserted into the variable region of the minimal synthetic promoter chassis (MinSyn000). D. Expression from MinSyns with combinations of AtTGA1 binding sites. Error bars indicate the mean and 2 x standard error of 3 replicates. Different lowercase letters indicate significant difference (ANOVA, *P* = 2.79e-11, TukeyHSD, α = 0.05).

## Methods

### Plasmid construction

Constructs were designed in Benchling (San Francisco, CA). TF coding sequences were synthesized (Twist Biosciences, San Francisco, CA) and HiBiT tag, stop codon (VSGWRLFKKIS*) and attB sites for Gateway cloning introduced by PCR. Amplicons were cloned into pDONOR207 using BP Clonase (ThermoFisher, Waltham, MA) and subcloned into pH9GW by LR-clonase (ThermoFisher). All other plasmids were assembled in BsaI-mediated Golden Gate reactions as previously described^6^. Details of all standard parts and assembled expression constructs are provided as Supporting Information. Plasmids and sequences are available at Addgene.

### Recombinant protein production

Bacteria harboring expression vectors were grown at 37°C, 220 rpm in 20 mL LB with (100 mg/L kanamycin) to OD_600_ 0.6-0.8. Expression was induced with 0.2 mM IPTG and cultures incubated at 18°C, 200 rpm for 18-20 h. Cells were collected in lysis buffer (50 mM Tris-HCL pH8.0, 500 mM NaCl, 20 mM Imidazole, 10% Glycerol, 0.05% Tween-20) and lysed by 20 cycles of sonication at 2s on, 5s off, 11 µS amplitude (Soniprep 150, MSE). Proteins were purified on 50 µL Ni-NTA resin and eluted in 100 µL elution buffer (50 mM Tris-HCL pH8.0, 500 mM NaCl, 300 mM Imidazole, 10% Glycerol, 0.05% Tween-20). Protein was quantified using the Nano-Glo® HiBiT Extracellular Detection System (N2420, Promega, Madison, WI) using the HiBiT Control Protein (20 µM, N3010, Promega) as a standard.

### EMSA

EMSA was performed as previously described ^6^ except that probes were produced by PCR using cy5 labeled oligonucleotide primers.

### qTFD assay

Double-stranded probes (DSPs) of random sequence or containing candidate binding sites were made by combining 2 µL of 100 µM forward and reverse oligos and 10 µL 2X annealing buffer (20 mM Tris-HCl pH8.0, 100 mM NaCl, 2 mM EDTA) in a 20 µL reaction. After heating to 95°C, the temperature was reduced by 0.1°C/s to 25°C. 5 µL 40 µg/µL of each DSP was combined with 45 µL DNA coating solution (17250, ThermoFisher Scientific) and added to six wells of a medium-binding microplate (655076, Greiner Bio-One, Kremsmünster, Germany) and incubated for 20 hrs at room temperature in the dark. Unbound probe was removed by three washes with 1X PBS. 270 µL of 3% bovine serum albumin (BSA) in PBS was added to three wells per probe and incubated at room temperature for 30 mins. After one wash with 1X PBS, 1.5 - 50 nM purified protein in binding buffer (25mM Tris-HCL pH8.0, 100 mM KCl, 2 mM DTT, 1 mM EDTA, 0.1 % BSA, 500 ng Poly(dI-dC), 5% Glycerol, 0.05% IGEPAL CA630) was added and plates incubated for 1-3 h at room temperature.

Unbound protein was removed by three washes of wash buffer (20 mM Tris-HCL pH8.0, 100 mM KCl, 10% Glycerol, 0.01% IGEPAL CA630). Bound protein was quantified using the Nano-Glo® HiBiT Extracellular Detection System (N2420, Promega). To quantify immobilized DNA in wells without protein, 0.25 µL PicoGreen (P7581, ThermoFisher Scientific) in 50 µL 1x TE buffer (10 mM Tris-HCL pH8.0, 1 mM EDTA) was incubated for 2 min at room temperature. Luminescence and fluorescence were quantified in a CLARIOstar Plus plate reader (BMG LABTECH, Ortenberg). Sequences of probes and example data and analysis are provided as Supporting Information.

### Protoplast preparation and transfection

Mesophyll protoplasts were prepared from *A. thaliana* leaves and transfected as previously described^3^. To quantify expression from MinSyns, 10 μg plasmid DNA comprising equal molar ratios of plasmids encoding MinSyn:LucF:*AtuOCSt* and a calibrator plasmid (*AtuNOSp*:LucN:*AtuNOSt*) was used to transfect 200 µL protoplasts (10^4^ -10^5^ /mL) and expression was normalized to an experiment calibrator (*AtuMASp*:LucF:*AtuOCSt* + *AtuNOSp*:LucN:*AtuNOSt*) as previously described^3^.

## Supporting Information

Details and sequences of plasmids and DNA probes are provided as supporting information files.

## Acknowledgements

pH9GW was a gift from Paul O’Maille. We thank Will Nash, Wilfried Haerty, Eftychios Frangedakis, Susana Sauret-Gueto, Tufan Oz, Siobhan Brady and Gozde Demirer for helpful discussions during the development of qTFD. We thank the John Innes Centre Horticultural services team for help with plant husbandry.

## Author contributions

YC and NP conceptualized the study. YC and SW analyzed sequences and produced constructs and protein. YC performed TF-binding and expression assays. YC and NP drafted the manuscript and all authors edited and approved.

