## Supporting Information for "Tuning plant promoters using a simple split luciferase method for quantifying transcription factor binding affinity"

Cai, Y-M[1], Witham, S[1], and Patron NJ[1]

#### Supporting Information

#### 1. Plasmids used in this study

| Plasmids for transcription factor protein expression | | | | | |
|---|---|---|---|---|---|
| **Addgene#** | **Plasmid code** | **Description** | **Acceptor** | **Plasmid type** | **Source of plasmid** |
| 196141 | pEPYCeGM0001 | AtTGA1:HiBiT | pDONR207 | Gateway Entry | This study |
| 196143 | pEPYCeGM0009 | AtNLP6:HiBiT | pDONR207 | Gateway Entry | This study |
| 196144 | pEPYCeGM0010 | AtNLP7:HiBiT | pDONR207 | Gateway Entry | This study |
| 196146 | pEPYCeGM0012 | AtNAC032:HiBiT | pDONR207 | Gateway Entry | This study |
| 196147 | pEPYCeGM0022 | AtARF18:HiBiT | pDONR207 | Gateway Entry | This study |
| 196148 | pEPYCdKN0001 | T7_9xHis:AtTGA1:HiBiT | pH9GW | Gateway Expression | This study |
| 196150 | pEPYCdKN0009 | T7_9xHis:AtNLP6:HiBiT | pH9GW | Gateway Expression | This study |
| 196151 | pEPYCdKN0010 | T7_9xHis:AtNLP7:HiBiT | pH9GW | Gateway Expression | This study |
| 196153 | pEPYCdKN0012 | T7_9xHis:AtNAC032:HiBiT | pH9GW | Gateway Expression | This study |
| 196154 | pEPYCdKN0022 | T7_9xHis:AtARF18:HiBiT | pH9GW | Gateway Expression | This study |
| | pH9GW | pDESTINATION LacI+T7_9xHIS | n/a | Gateway Destination | Paul O'Maille |

| Level 0 Phytobricks | | | | | | | |
|---|---|---|---|---|---|---|---|
| **Addgene#** | **Plasmid code** | **Part type** | **Description** | **Compatibility with Assembly Systems** | **Cloning overhang (top strand)** | | **Source of plasmid** |
| | | | | | **5'** | **3'** | |
| 196155 | pEPYC0CM0515 | PROM | MinSyn_301 | MoClo, Loop, GB | GGAG | TACT | This study |
| 196156 | pEPYC0CM0519 | PROM | MinSyn_302 | MoClo, Loop, GB | GGAG | TACT | This study |
| 196157 | pEPYC0CM0520 | PROM | MinSyn_303 | MoClo, Loop, GB | GGAG | TACT | This study |
| 196158 | pEPYC0CM0522 | PROM | MinSyn_305 | MoClo, Loop, GB | GGAG | TACT | This study |
| 196159 | pEPYC0CM0513 | PROM | MinSyn_308 | MoClo, Loop, GB | GGAG | TACT | This study |
| 196160 | pEPYC0CM0528 | PROM | MinSyn_310 | MoClo, Loop, GB | GGAG | TACT | This study |
| 196161 | pEPYC0CM0529 | PROM | MinSyn_311 | MoClo, Loop, GB | GGAG | TACT | This study |
| 154503 | pEPYC0CM0035 | PROM | MinSyn_000 | MoClo, Loop, GB | GGAG | TACT | Cai et al., 2020 |
| 50255 | pICH42211 | PROM | AtuNOSp | MoClo, Loop, GB | GGAG | TACT | Engler et al., 2014 |
| 50272 | pICH85281 | PROM | AtuMASp | MoClo, Loop, GB | GGAG | TACT | Engler et al., 2014 |
| 50285 | pICH41402 | 5UTR | TMVΩ | MoClo, Loop, GB | TACT | AATG | Engler et al., 2014 |
| 154594 | pEPAS0CM0008 | CDS | LucF | MoClo, Loop, GB | AATG | TTCG | Cai et al., 2020 |
| 154595 | pEPYC0CM0133 | CDS | LucN | MoClo, Loop, GB | AATG | TTCG | Cai et al., 2020 |
| 50308 | pICSL50007 | CTAG | FLAG tag | MoClo, Loop, GB | TTCG | GCTT | Engler et al., 2014 |
| 50343 | pICH41432 | 3UTR+TERM | AtuOCSt | MoClo, Loop, GB | GCTT | CGCT | Engler et al., 2014 |
| 50339 | pICH41421 | 3UTR+TERM | AtuNOSt | MoClo, Loop, GB | GCTT | CGCT | Engler et al., 2014 |

| Level 1 Assemblies | | | | | |
|---|---|---|---|---|---|
| **Addgene#** | **Plasmid code** | **Plasmid type** | **Contents** | **Acceptor** | **Source of plasmid** |
| 196166 | pEPYC1CB0597 | Plant expression (promoter:LucF) | MinSyn_301 (pEPYC0CM0515)_TMV(pICH41402)_LucF (pEPAS0CM0008)_FLAG(pICSL50007)_ocsT(pICH41432) | pICH47732 (Addgene 48000) | This study |
| 196167 | pEPYC1CB0601 | Plant expression (promoter:LucF) | MinSyn_302 (pEPYC0CM0519)_TMV(pICH41402)_LucF (pEPAS0CM0008)_FLAG(pICSL50007)_ocsT(pICH41432) | pICH47732 (Addgene 48000) | This study |
| 196168 | pEPYC1CB0602 | Plant expression (promoter:LucF) | MinSyn_303 (pEPYC0CM0520)_TMV(pICH41402)_LucF (pEPAS0CM0008)_FLAG(pICSL50007)_ocsT(pICH41432) | pICH47732 (Addgene 48000) | This study |
| 196169 | pEPYC1CB0604 | Plant expression (promoter:LucF) | MinSyn_305 (pEPYC0CM0522)_TMV(pICH41402)_LucF (pEPAS0CM0008)_FLAG(pICSL50007)_ocsT(pICH41432) | pICH47732 (Addgene 48000) | This study |
| 196170 | pEPYC1CB0595 | Plant expression (promoter:LucF) | MinSyn_308 (pEPYC0CM0513)_TMV(pICH41402)_LucF (pEPAS0CM0008)_FLAG(pICSL50007)_ocsT(pICH41432) | pICH47732 (Addgene 48000) | This study |
| 196171 | pEPYC1CB0607 | Plant expression (promoter:LucF) | MinSyn_310 (pEPYC0CM0528)_TMV(pICH41402)_LucF (pEPAS0CM0008)_FLAG(pICSL50007)_ocsT(pICH41432) | pICH47732 (Addgene 48000) | This study |
| 196172 | pEPYC1CB0608 | Plant expression (promoter:LucF) | MinSyn_311 (pEPYC0CM0529)_TMV(pICH41402)_LucF (pEPAS0CM0008)_FLAG(pICSL50007)_ocsT(pICH41432) | pICH47732 (Addgene 48000) | This study |
| 154630 | pEPYC1CB0007 | Plant expression (promoter:LucF) | MinSyn_000 (pEPYC0CM0035)_TMV(pICH41402)_LucF (pEPAS0CM0008)_FLAG(pICSL50007)_ocsT(pICH41432) | pICH47732 (Addgene 48000) | Cai et al., 2020 |
| 154654 | pEPYC1CB0197 | Plant expression calibrator (promoter:LucN) | NOSp(pICH42211)_TMV(pICH41402)_LucN (pEPAS0CM0133)_FLAG(pICSL50007)_nosT(pICH41421) | pICH47732 (Addgene 48000) | Cai et al., 2020 |
| 154655 | pEPYC1CB0199 | Plant expression experiment calibrator (promoter:LucF) | MASp(pICH85281)_TMV(pICH41402)_LucF (pEPAS0CM0008)_FLAG(pICSL50007)_ocsT(pICH41432) (experiment normaliser) | pICH47732 (Addgene 48000) | Cai et al., 2020 |

#### 2. Sequences of DNA probes

| Name and source | TF | Oligo ID | Sequence |
|---|---|---|---|
| Random (rnd) | NONE | ptoz179_PS_NC_F | TAGCGAAGTACGATCCCATGAAGACGCTGGGTTTACATGGAATGGTGCTTCTGTTCTAACAGGCTAGGATATAAGGCCATCACGCAGTA |
| | | ptoz180_PS_NC_R | TACTGCGTGATGGCCTTATATCCTAGCCTGTTAGAACAGAAGCACCATTCCCATGTAAACCCAGCGTCTTCATGGGATCGTACTTCGCTA |
| CaMV35s | TGA1 | YMC175 | ATGAAGACGCTGGGTTTACATGGGAATGGCTGACGTAAGGGATGACGCACATGCTTCTGTTCTAACAGGCTAGGATATAA |
| | | YMC176 | TTATATCCTAGCCTGTTAGAACAGAAGCATGTGCGTCATCCCTTACGTCAGCCATTCCCATGTAAACCCAGCGTCTTCAT |
| NIR1 | NLP6/ NLP7 | ptoz177_PS_PC_F | TAGCGAAGTACGATCCCATCAAAGAGAAACAACTTGACCCTTTACATTGCTCAAGAGCTCATCTCTTCCCTCTACGGCCATCACGCAGTA |
| | | ptoz178_PS_PC_R | TACTGCGTGATGGCCGTAGAGGGAAGAGATGAGCTCTTGAGCAATGTAAAGGGTCAAGTTGTTTCTCTTTGATGGGATCGTACTTCGCTA |
| ANAC032SO1 | ANAC032 | ptoz369_PC2_ANAC032_F | TAGCGAAGTACGATCCCATGAAGACGGAGGTAAGCAAATTGATCACGCAACTGGTGGATATAAGGCCATCACGCAGTA |
| | | ptoz370_PC2_ANAC032_R | TACTGCGTGATGGCCTTATATCCACCAGTTGCGTGATCAATTTGCTTACCTCCGTCTTCATGGGATCGTACTTCGCTA |
| DR5(7X) | ARF18 | DR5F | CCTTTTGTCTCCCTTTTGTCTCCCTTTTGTCTCCCTTTTGTCTCCCTTTTGTCTCCCTTTTGTCTCCCTTTTGTCTC |
| | | DR5R | GAGACAAAAGGGAGACAAAAGGGAGACAAAAGGGAGACAAAAGGGAGACAAAGGGAGACAAAAGGGAGACAAAAGG |
| TFBS_07 AT3G14060 | TGA1 | YMC183 | TCCCATGAAGACGCTGGGTTTACATGGGAATGCATGACATCAACGTGGTGCTTCTGTTCTAACAGGCTAGGATATAAGGC |
| | | YMC184 | GCCTTATATCCTAGCCTGTTAGAACAGAAGCACCACGTTGATGTCATGCATTCCCATGTAAACCCAGCGTCTTCATGGGA |
| TFBS_05 AT3G14205 | TGA1 | YMC187 | TCCCATGAAGACGCTGGGTTTACATGGGAATGCATTACATCATCATAGTGCTTCTGTTCTAACAGGCTAGGATATAAGGC |
| | | YMC188 | GCCTTATATCCTAGCCTGTTAGAACAGAAGCACTATGATGATGTAATGCATTCCCATGTAAACCCAGCGTCTTCATGGGA |
| TFBS_09 AT1G68490 | TGA1 | YMC189 | TCCCATGAAGACGCTGGGTTTACATGGGAATGTGTCACATCAGCATAGTGCTTCTGTTCTAACAGGCTAGGATATAAGGC |
| | | YMC190 | GCCTTATATCCTAGCCTGTTAGAACAGAAGCACTATGCTGATGTGACACATTCCCATGTAAACCCAGCGTCTTCATGGGA |
| TFBS_02 AT1G77450 | TGA1 | ptoz329_PS_ANAC032_3F | TAGCGAAGTACGATCCCGGACCGCTACATTCCAAATAGTCTGACGTAAGCAATGACAAAACTCACCTACATGGCCATCACGCAGTA |
| | | ptoz330_PS_ANAC032_3R | TACTGCGTGATGGCCATGTAGGTGAGTTTTGTCATTGCTTACGTCAGACTATTTGGAATGTAGCGGTCCGGGATCGTACTTCGCTA |
| TFBS_12 AT1G64530 | TGA1 | ptoz307_PS_NLP6_2F | TAGCGAAGTACGATCCCAACACGCACTTAGTGCCTAATTACGTCTTAATTTAATATTTTTTAAAGCGGCCATCACGCAGTA |
| | | ptoz308_PS_NLP6_2R | TACTGCGTGATGGCCGCTTTAAAAAATATTAAATTAAGACGTAATTAGGCACTAAGTGCGTGTTGGGATCGTACTTCGCTA |
| TFBS_14 AT3G61830 | TGA1 | ptoz555_Pr_104_PS_ARF18_3F | TAGCGAAGTACGATCCCTGAGTTTCCTTTTTAGCGATGACATGATAAAACAAAAAACAACAATTTGGCCATCACGCAGTA |
| | | ptoz556_Pr_104_PS_ARF18_3R | TACTGCGTGATGGCCAAATTGTTGTTTTTTGTTTTATCATGTCATCGCTAAAAAGGAAACTCAGGGATCGTACTTCGCTA |

### 3. Example data and analysis

| | 0 nM | 1.5 fM | 5 nM | 15 nM | 2x free probe 15 nM | 5x free probe 15 nM protein | 10x free probe 15 nM protein | rnd probe 15 nM protein |
|---|---|---|---|---|---|---|---|---|
| HiBit Rep1 | 579 | 224535 | 567146 | 804307 | 49390 | 28536 | 23612 | 324075 |
| HiBit Rep2 | 600 | 248884 | 560068 | 940370 | 63746 | 42783 | 26734 | 397371 |
| HiBit Rep3 | 421 | 306935 | 793533 | 1192230 | 119946 | 39637 | 35014 | 398011 |
| Pico Rep 1 | 16859 | 16859 | 16859 | 16859 | 16859 | 16859 | 16859 | 20810 |
| Pico Rep 2 | 9102 | 9102 | 9102 | 9102 | 9102 | 9102 | 9102 | 15166 |
| Pico Rep 3 | 12382 | 12382 | 12382 | 12382 | 12382 | 12382 | 12382 | 9305 |
| mean | 12781 | 12781 | 12781 | 12781 | 12781 | 12781 | 12781 | 15093.66667 |
| HiBit/Pico Rep1 | 0.04530162 | 17.5678742 | 44.3741491 | 62.9298959 | 3.864329865 | 2.232689148 | 1.847429779 | 21.470926 |
| HiBit/Pico Rep2 | 0.04694468 | 19.4729677 | 43.8203583 | 73.5756201 | 4.987559659 | 3.347390658 | 2.091698615 | 26.3270025 |
| HiBit/Pico Rep3 | 0.03293952 | 24.0149441 | 62.0869259 | 93.2814334 | 9.384711681 | 3.101244034 | 2.739535248 | 26.36940439 |
| mean | 0.04172861 | 20.3519286 | 50.0938111 | 76.5956498 | 6.078867068 | 2.893774613 | 2.226221214 | 24.72244429 |
| 2xSE | 0.00884013 | 3.8245883 | 11.997376 | 17.7818031 | 3.36885099 | 0.676187878 | 0.532334843 | 3.251610454 |
| Tukey HSD | d | c | b | a | cd | cd | cd | c |

| | TGA probe | Rnd Probe |
|---|---|---|
| **Rep1 normalised to rnd (rQ)** | 2.54545607 | 0.868479093 |
| **Rep2 Normalised to rnd (rQ)** | 2.97606576 | 1.064902895 |
| **Rep3 Normalised to rnd (rQ)** | 3.77314768 | 1.066618012 |
| **mean** | 3.09822317 | 1 |
| **2xSE** | 0.71925748 | 0.131524635 |

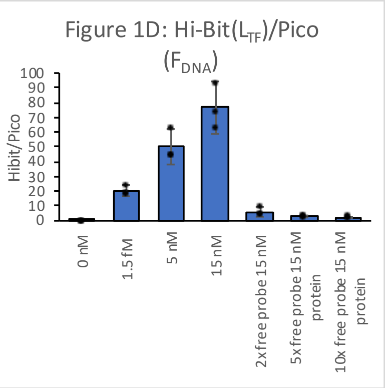

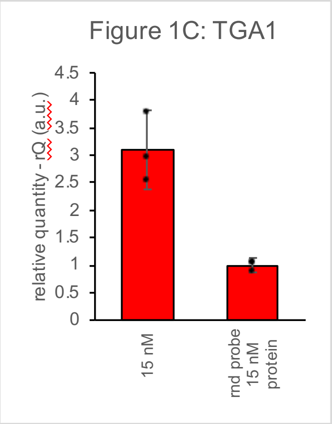
